# Resistance to PSEN1-selective γ-secretase inhibitors in T-cell acute lymphoblastic leukemia

**DOI:** 10.1101/2024.03.01.582944

**Authors:** Charlien Vandersmissen, Sofie Demeyer, Kris Jacobs, Lien Boogaerts, Sara Gutiérrez Fernández, Heidi Segers, Lucía Chávez-Gutiérrez, Jan Cools

## Abstract

PSEN1-selective gamma-secretase inhibitors (GSI), such as MRK-560, are a potential option for the treatment of T-cell acute lymphoblastic leukemia (T-ALL) with NOTCH1 activating mutations, as these show less toxicity compared to broad-spectrum GSIs. However, an important challenge with targeted therapies for cancer treatment is the rapid development of drug resistance. We therefore investigated if *PSEN1* mutations could confer resistance to MRK-560 in T-ALL. We performed a CRISPR-mediated mutagenesis screen in a T-ALL cell line to identify mutations leading to MRK-560 resistance and confirmed these findings in additional cell lines. We identified 3 types of resistance mutations. Mutations at the enzyme-drug interface directly disrupt the interaction of MRK-560 with PSEN1. Mutations at the enzyme-substrate interface cause a shift in relative binding affinities towards drug and/or substrate. The third resistance mechanism involves a mutation at the enzyme-substrate interface that hinders the entrance of MRK-560 to the binding pocket. These findings contribute to the understanding of the PSEN1-selectivity of MRK-560 and can help to design other PSEN1-selective GSIs to overcome resistance in cancer therapy.

## Introduction

The NOTCH signaling pathway is involved in multiple mechanisms in the human body, such as embryonic development, cell proliferation, differentiation, apoptosis and cell fate decisions.^1,2^ The NOTCH proteins are transmembrane receptors that are activated by interaction with their ligands (Delta-like or Jagged ligands), expressed on the surface of adjacent cells.^3^ This interaction results in conformational changes in the receptor and proteolytic cleavage by the ADAM10/ADAM17/ADAMTS1 metalloprotease, followed by a second cleavage event by the γ-secretase complex.^1,2^ As a result of these sequential cleavage events, the intracellular domain of the NOTCH receptor (NICD) is released from the membrane and translocates to the nucleus, where it functions as a transcriptional regulator of its target genes, such as the HES/HEY gene family. The NOTCH activity is further regulated by the F-box/WD repeat-containing protein 7 (FBXW7), an E3 ubiquitin ligase, which recognizes the C-terminal PEST domain of the NICD and targets NICD for proteasomal degradation, resulting in the termination of NOTCH signaling.^1,2^

Mammals have four NOTCH receptors (NOTCH1, NOTCH2, NOTCH3, NOTCH4) with slightly different structures/functions and distinct subcellular localizations.^1^ While all NOTCH receptors play some role in cancer development, this is most clear for NOTCH1, which is frequently mutated or overexpressed in breast cancer^4^, lung cancer^5^, aggressive desmoid tumors^6^ and in hematological cancers, such as T-cell acute lymphoblastic leukemia (T-ALL)^7^, chronic lymphocytic leukemia (CLL)^8^, diffuse large B-cell lymphoma (DLBCL)^9^ and mantle cell lymphoma (MCL)^10^. The majority of T-ALL cases harbor activating NOTCH1 mutations and therefore NOTCH1 has been actively investigated as a major target for therapy in this rare leukemia.^11–13^

One way to inhibit NOTCH1 signaling is by treatment with γ-secretase inhibitors (GSI). Indeed, both overexpressed and mutant NOTCH1 in cancer cells remain dependent on proteolytic cleavage by the γ-secretase complex. The γ-secretase complex consists of four different proteins: nicastrin (NCSTN), anterior pharynx-defective 1 (APH1A/B), presenilin (PSEN1/2) and presenilin enhancer 2 (PSENEN, also known as PEN-2).^14,15^ There are 4 different subtypes of the γ-secretase complex with APH1A or APH1B included and with PSEN1 or PSEN2 included. Presenilin contains the active site and is responsible for the cleavage of multiple substrates, including NOTCH1 and the amyloid precursor protein (APP). In the past, broad-spectrum GSIs that target both PSEN1 and PSEN2 γ-secretase complexes were developed and showed potent activity against T-ALL cell lines.^16^ Unfortunately, clinical trials in different cancer types with overactive NOTCH1 signaling were negative.^17,18^ Patients treated with broad-spectrum GSI suffered from dose-limiting gastro-intestinal toxicities and consequently interest in these inhibitors decreased.^18^ Interestingly, one broad-spectrum GSI, namely nirogacestat, was recently approved by the Food and Drug administration for desmoid tumors, demonstrating the clinical potential.^19^

One possibility to lower the toxicity of GSIs is by more precisely targeting of a subset of γ-secretase complexes.^20^ For example, Habets *et al.* discovered that T-ALL cells predominantly contain PSEN1 γ-secretase complexes, whereas other cell types, like gastro-intestinal cells, express both PSEN1 and PSEN2 γ-secretase complexes.^21^ Moreover, they showed that PSEN1 selective γ-secretase inhibition with the drug MRK-560 resulted in effective NOTCH1 inhibition without gastro-intestinal toxicities in mice. Additional preclinical experiments have shown that MRK-560 can be used to safely target T-ALL in cell lines and patient-derived xenograft mouse models.^22,23^

These findings open new avenues for the development of PSEN1-selective inhibitors for the treatment of T-ALL, but several questions remain to be addressed. One potential limitation with the use of targeted treatments for cancer therapy is the development of resistance. For instance, *PTEN* inactivation is known to confer resistance to NOTCH1 inhibition via upregulation of PI3K-AKT signaling pathway.^24^ However, it is still unknow if mutations in *PSEN1* could give resistance towards PSEN1-selective GSI in T-ALL. Therefore, we performed CRISPR-Cas9 genome editing in a T-ALL cell line to introduce random mutations in *PSEN1* and to subsequently screen for resistance mutations after MRK-560 treatment. Identification of these resistance-associated mutations will enhance our understanding of PSEN1-selectivity of MRK-560, facilitating the design of other PSEN1-selective γ-secretase inhibitors to target NOTCH1 activation and overcome resistance.

## Materials and methods

### Cell lines and compounds

RPMI-8402, DND-41 and HPB-ALL cells were obtained from DSMZ, and cell identification was confirmed by sequencing of *NOTCH1* mutations and by short tandem repeat (STR) analysis. These cells were cultured in RPMI-1640 medium supplemented with 20% fetal bovine serum (FBS, Gibco) and were transduced with Cas9 to create stable Cas9-expressing cell lines. *Psen1*^-/-^ *Psen2*^-/-^ mouse embryonic fibroblasts (MEFs) and HEK293T/17 cells were cultured in DMEM/F-12 (Fisher scientific) with 10% FBS. MRK-560 was purchased from Tocris Bioscience (#4000); crenigacestat (LY3039478, #HY-12449) and nirogacestat (PF-03084014, #HY-15185) were purchased from MedChem Express.

### CRISPR screening in RPMI-8402 Cas9 cells

We designed sgRNAs against the coding exons of human *PSEN1* (Exon 3 – Exon 12) with Benchling and we ordered the sgRNAs as single-stranded oligos at Integrated DNA Technologies (IDT). After annealing and pooling of the guides of one exon, we introduced these gRNA oligos in a retroviral pMx-U6-gRNA-SFFV-mCherry vector. Next, we transfected HEK293T cells with the pooled pMx vector library of one exon, together with gag-pol plasmid (#14887, addgene) and VSVG plasmid (#8454, addgene) and the produced VSVG-pseudotyped retroviral vectors were harvested after 48 hours. We transduced RPMI-8402 cas9 cells on retronectine-coated plates with the VSVG-pseudotyped retroviral vectors via spinfection and obtained transduction efficiencies below 30%. The cells were washed 24h after spinfection and were sorted based on mCherry expression the day afterwards. The cells transduced with the sgRNA pools targeting different exons of *PSEN1* were treated with 0.1 μM MRK-560 until resistance cells were observed. We increased the dose of MRK-560 every 2 weeks up to a final concentration of 20 μM and collected DNA and protein lysates at each different concentration.

### Validation of *PSEN1* resistance mutations in MEFs

Initially, we established a stable *Psen1*^-/-^ *Psen2*^-/-^ MEF cell line expressing a truncated form of human NOTCH1 (ΔEGF-L1600P-ΔPEST). To achieve this, HEK293T/17 cells were transfected with a retroviral construct containing human mutant NOTCH1 (ΔEGF-L1600P-ΔPEST)-IRES-GFP in combination with the PIK helper plasmid, and the resulting retroviral vectors were used to transduce *Psen1*^−/−^ *Psen2*^−/−^ MEF cells and GFP-expressing cells were selected.^25^ These cells were transduced with viral vectors containing mutated human *PSEN1*. Specific mutations in human *PSEN1* were obtained by Q5 site-directed mutagenesis of the pMSCV-puro vector containing wild type human *PSEN1.*^26^ We designed primers for Q5 site-directed mutagenesis (New England Biolabs) with NEBaseChanger.neb.com and performed site-directed mutagenesis according to the manufacturer’s protocol. In this way, we obtained stable hNOTCH1 expression MEFs with either the *hPSEN1* wild type gene or a mutated version. We treated these different MEF cell lines with DMSO, MRK-560 or crenigacestat for 24 or 48 hours and collected protein lysates.

### Determination of Aβ production by ELISA

MEF cells with WT or mutant PSEN1 were first transduced with adeno-associated virus containing the C-terminal part of APP (APP-C99). Two days after transduction, the conditioned media containing secreted Aβ fragments was collected. Aβ37, Aβ38, Aβ40 and Aβ42 product levels were quantified on Multi-spot 96-wel plates precoated with anti-Aβ37, anti-Aβ38, anti-Aβ40 and anti-Aβ42 antibodies obtained from Janssen Pharmaceutica using multiplex MSD technology. MSD plates were blocked with 0.1% casein/PBS for 2h and washed before adding the samples. The detection antibody (SULFO-TAG JRF/AbN/25) was diluted in blocking buffer and was mixed with the standards (Aβ37, Aβ38, Aβ40 and Aβ42 peptides) or reaction samples and these mixtures were loaded on the MSD plates. After overnight incubation, plates were rinsed and MSD read buffer T was added to read out the plates with MSD Sector Imager 6000.

### Generation of specific *PSEN1* mutations in the genome of T-ALL cell lines

We designed sgRNAs and homology direct repair (HDR) oligos to introduce three different mutations (275(A>Y), 275(A>Y)+276(Q>E) and 421_422(->ILS)) in T-ALL cell lines. A list of the used sgRNAs and HDR oligos is given in **Table 1**. We electroporated three Cas9-expressing T-ALL cell lines (RPMI-8402, DND-41, HPB-ALL) with the combination of a sgRNA and HDR oligo using the MPK5000 nucleofector from Thermofisher. Next, we cultured the cells in the presence of 1 μM MRK-560 and after treatment of 3 weeks, we determined if the majority of the cells harbored the correct mutation in *hPSEN1* by Sanger sequencing. In this way, we were able to introduce each mutation separately in three T-ALL cell lines. These cells were treated with DMSO, 1 μM MRK-560, 5 μM MRK-560, 1 μM crenigacestat or 1 μM nirogacestat for 2 days to isolate protein lysates or for 10 days to follow cell growth.

**Table 1.**
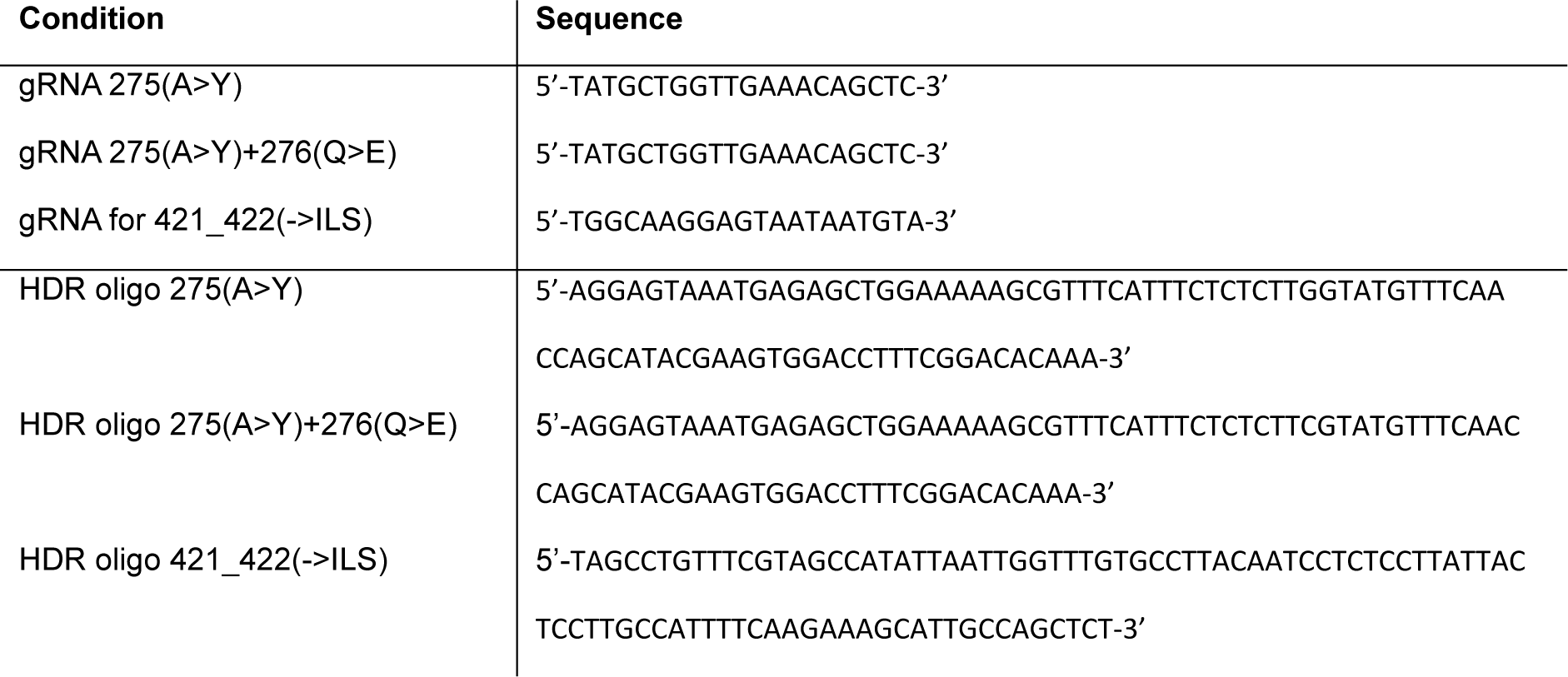
Overview of the used sgRNAs and HDR oligos for genome editing and generating specific mutations in T-ALL cells.

### Bulk DNA sequencing

Exon-specific primers were designed for exon 3 to 12 of human *PSEN1*. PCR was performed with 2x KAPA HiFi hotstart readymix, which was followed with PCR clean-up using AMP pure beads. A second KAPA HiFi hotstart PCR was performed with illumina index primers, followed by purification via AMP pure beads. These PCR products were sequenced with an Illumina MiSeq sequencer. The resulting paired-end sequencing reads were cleaned with fastq-mcf and quality control was performed with FastQC. The high-quality reads were then mapped to the human reference genome (hg38) with HISAT2. Further processing was performed with SAMtools. The mutations were then called and filtered with BCFtools, where we limited our area of investigation to the exons that were targeted.

### Western blotting

Protein lysates were obtained using 1X Cell Lysis Buffer (#9803, Cell Signaling), which contained 5mM Na_3_VO_4_ (#P0758S, New England Biolabs) and a protease inhibitor (cOmplete, EDTA-free, Sigma) after the indicated treatment. Afterwards, proteins were separated by SDS-PAGE (NuPAGE NOVEX 4–12% Bis Tris, Invitrogen) and transferred to a PVDF membrane using the Mini Trans-Blot Cell system (Bio-rad). Primary antibodies for full NOTCH1 (#3268, Cell Signaling), cleaved NOTCH1 V1744 (#4147, Cell signaling), HES1 (#11988, Cell Signaling), PSEN1 NTF (#87146, Cell Signaling), PSEN1 CTF (#5643, Cell signaling), PEN-2 (#8598, Cell signaling), nicastrin (9C3, kindly provided by Prof. Wim Annaert)^27^ or ß-actin (#A1978, Sigma), were mixed in blocking buffer and applied to the blot overnight (4°C). Detection was carried out with secondary antibodies conjugated with horse-radish peroxidase. Images were obtained using Vilber Fusion FX6 imager and quantification of the bands was performed with the Vilber Fusion software.

### qPCR

RNA was extracted with the Maxwell RSC instrument using the RSC simplyRNA Cells Kit (#AS1390, Promega) after the indicated treatment. Subsequent cDNA synthesis was performed with GoScript Reverse Transcriptase kit (Promega). Finally, qPCR was conducted with GoTaq qPCR master mix (Promega) for the desired genes and housekeeping genes of which the primer sequences are given in **Table 2**. Fold changes of mRNA levels were calculated with the delta-delta-Ct method.

**Table 2.**
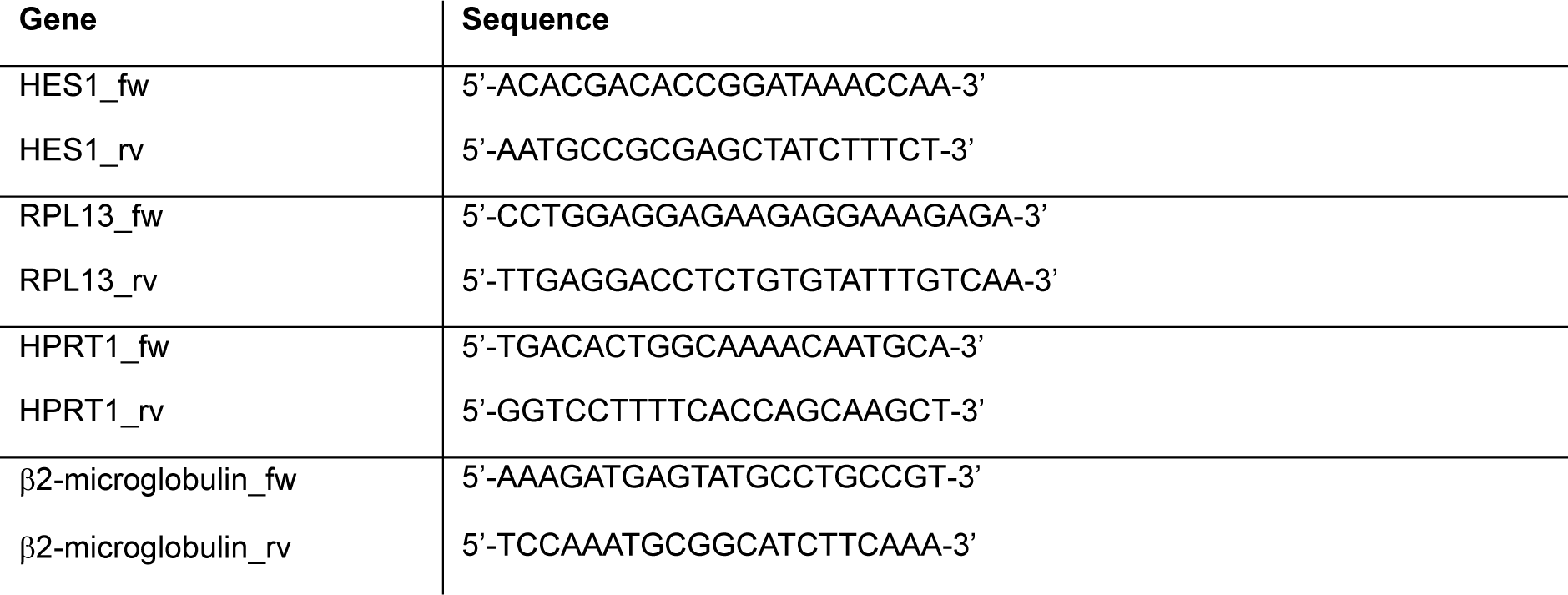
qPCR primers for human housekeeping genes and HES1.

## Results

### CRISPR-Cas9 mediated random mutagenesis of PSEN1 confers resistance to MRK-560 in T-ALL

We have previously shown that T-ALL cell lines with NOTCH1 mutations are sensitive to GSIs.^22,23^ Similar to primary T-ALL leukemia cells, these cell lines only express *PSEN1 (*and not *PSEN2)* and are sensitive to broad-spectrum GSIs as well as to PSEN1-selective GSIs, such as MRK-560.^21,22^

To identify if mutations in the human *PSEN1* gene could confer resistance to MRK-560, we used CRISPR-Cas9 genome editing to introduce random mutations in *PSEN1* in the T-ALL cell line RPMI-8402. This cell line has a mutation in the heterodimerization domain of the NOTCH1 receptor and is known to be dependent on the NOTCH1 signaling pathway for proliferation and survival. As a consequence, these cells are sensitive to GSI, including MRK-560. The RPMI-8402 cells were transduced with a lentiviral vector expressing Cas9 and we confirmed that these cells were still sensitive to MRK-560.

To introduce random mutations in the various coding exons of *PSEN1*, we designed 374 sgRNAs targeting exons 3 to 12 of *hPSEN1* (**Fig. 1A**) and we pooled the sgRNAs per exon to generate 10 pools and an 11^th^ pool of control sgRNAs. We cloned the sgRNA pools in a retroviral vector also expressing mCherry and transduced RPMI-8402 Cas9 cells with each of the 11 pools. The cells were selected based on expression of the mCherry fluorescent marker and were subsequently treated with increasing doses of MRK-560 over a period of approximately 18 weeks. The concentration of MRK-560 was initially 100 nM and was increased every 2 weeks to a final concentration of 20 µM.

**Figure 1.**
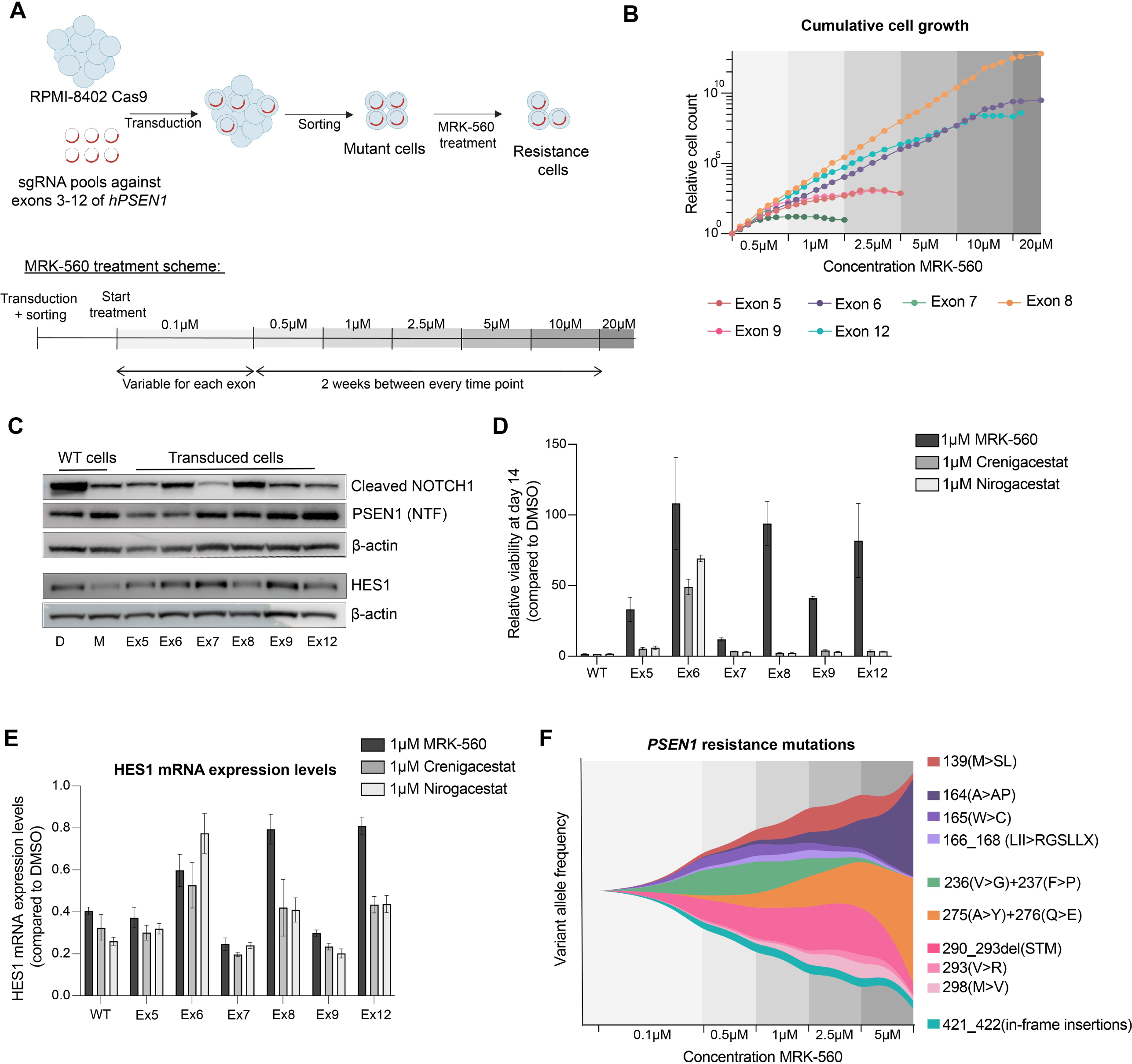
CRISPR-Cas9 mediated mutagenesis of *PSEN1* induces resistance to MRK-560 in a human T-ALL cell line. **(A)** Experimental set-up of the CRISPR screening of RPMI-8402 cells. **(B)** Cumulative cell growth of cells transduced with sgRNA library against different exons of *PSEN1*. The cell counts were calculated relatively to the cell count at start of 0.5 μM MRK-560 treatment. **(C)** Protein levels of cleaved NOTCH1(V1744), HES1, PSEN1 NTF and β-actine for RPMI-8402 Cas9 wild type (WT) cells or for RPMI-8402 Cas9 cells transduced with the pools of sgRNAs treated for 24 hours with DMSO (D) or 0.1 μM MRK-560 (M). **(D)** Relative viability percentage of RPMI-8402 Cas9 cells WT or transduced cells treated for 14 days with DMSO, 1 μM MRK-560, 1 μM crenigacestat, 1 μM nirogacestat. **(E)** Relative mRNA expression level of HES1 of RPMI-8402 Cas9 WT or transduced cells treated for 24 hours with DMSO, 1 μM MRK-560, 1 μM crenigacestat or 1 μM nirogacestat. **(F)** Fish plot of variant allele frequencies of the different mutations (of the different exons) over time determined with illumina Miseq bulk DNA sequencing. NTF: N-terminal fragment. CTF: C-terminal fragment. In panel D and E mean and standard deviation (error bars) of 3 replicates are shown.

Cells transduced with sgRNA pools targeting exon 5, 6, 7, 8, 9 or 12 were rapidly showing resistance to MRK-560 treatment. The most resistant cells were the cells transduced with sgRNAs targeting exon 6, 8 or 12 of *PSEN1*, which were characterized with high cell counts during MRK-560 treatment (**Fig. 1B**). After resistance to MRK-560 occurred, we detected the protein expression levels of cleaved NOTCH1, PSEN1 and HES1, a downstream NOTCH1 target, and we observed that the enhanced proliferation of these resistant populations was associated with sustained NOTCH1 signaling (**Fig. 1C**). In addition, we also treated these resistant cell populations with DMSO, MRK-560 and two broad-spectrum GSIs for 24h to detect HES1 mRNA levels (**Fig. 1E**), or for two weeks to evaluate growth inhibition (**Fig. 1D**). The growth curves showed that resistant cells transduced with sgRNA pools targeting exon 6, 8 and 12 were almost insensitive to MRK-560 treatment, since the relative viability was > 80% (**Fig. 1D, Fig. EV1**). Moreover, resistance to broad-spectrum GSIs was mainly observed for cells transduced with sgRNA pool against exon 6. The HES1 expression levels corresponded with the growth curves, showing high HES1 expression levels and consequently high NOTCH1 signaling for resistance cells with mutations in exon 6, 8 and 12 after treatment with MRK-560 (**Fig. 1E**).

To identify the specific resistant mutations in each exon, we amplified the various exons of *PSEN1* and subjected these to bulk DNA sequencing on an Illumina MiSeq run at each time point during MRK-560 treatment. This approach allowed us to track the different mutations over time and to identify the most resistant mutations (**Fig. 1F**). In total, we identified 12 potential resistance mutations, including point mutations, insertions and deletions. In exon 5, 7 and 8, we only found one genomic alteration in the resistance cells. In exon 5, we found an insertion of three nucleotides, resulting in the mutation 139(M>SL). Conversely, point mutations were discovered in exon 7 and 8, affecting two subsequent amino acids, respectively 236(V>G) + 237(F>P) and 275(A>Y) + 276(Q>E). Furthermore, we identified an in-frame insertion of three nucleotides at 164(A>AP), a point mutation 165(W>C) and a frameshift variant 166_168(LII>RGSLLX) in resistance cells transduced with sgRNA pool against exon 6. We also found three different alterations in exon 9: a deletion of 9 nucleotides 290_292del(STM), 293(V>R) and 298(M>V). Lastly, in exon 12, we identified three different in -frame insertions between amino acids 421 and 422: 421_422(->A), 421_422(->E) and 421_422(->ILS). All mutations are shown in **Fig. 1F**. Based on the frequency of the mutations in the next-generation sequencing data, the most resistant mutations seemed to be 164(A>AP), 275(A>Y) + 276(Q>E) and a deletion at amino acids 290-292.

### Confirmation of PSEN1 resistance mutations in mouse embryonic fibroblasts

To identify and confirm the true resistance mutations, we generated each of the mutants identified by next-generation sequencing in a PSEN1 expression construct and used mouse embryonic fibroblasts (MEF) deficient in *Psen1* and *Psen2* as model to validate our mutations. First, we transduced *Psen1*^−/−^ *Psen2*^−/−^ MEF cells with retroviral vector containing a construct with human NOTCH1(ΔEGF-L1600P-ΔPEST)-IRES-GFP to constitutively express mutant NOTCH1 (**Fig. 2A**). Next, we transduced these cells with retroviral vectors containing wild type or mutant human PSEN1. After selection with puromycin we were able to obtain stable MEFs with constitutively active NOTCH1 in combination with either wild type or mutated PSEN1. The presence of glycosylated (mature) nicastrin, PEN-2 and endoproteolyzed PSEN1 C-terminal fragments (CTF) indicated that all mutant PSEN1 were incorporated into the γ-secretase complex and rescued (at least partially) its activity, except for the frameshift mutation 166_168(LII>RGSLLX) (**Fig EV. 2**).

**Figure 2.**
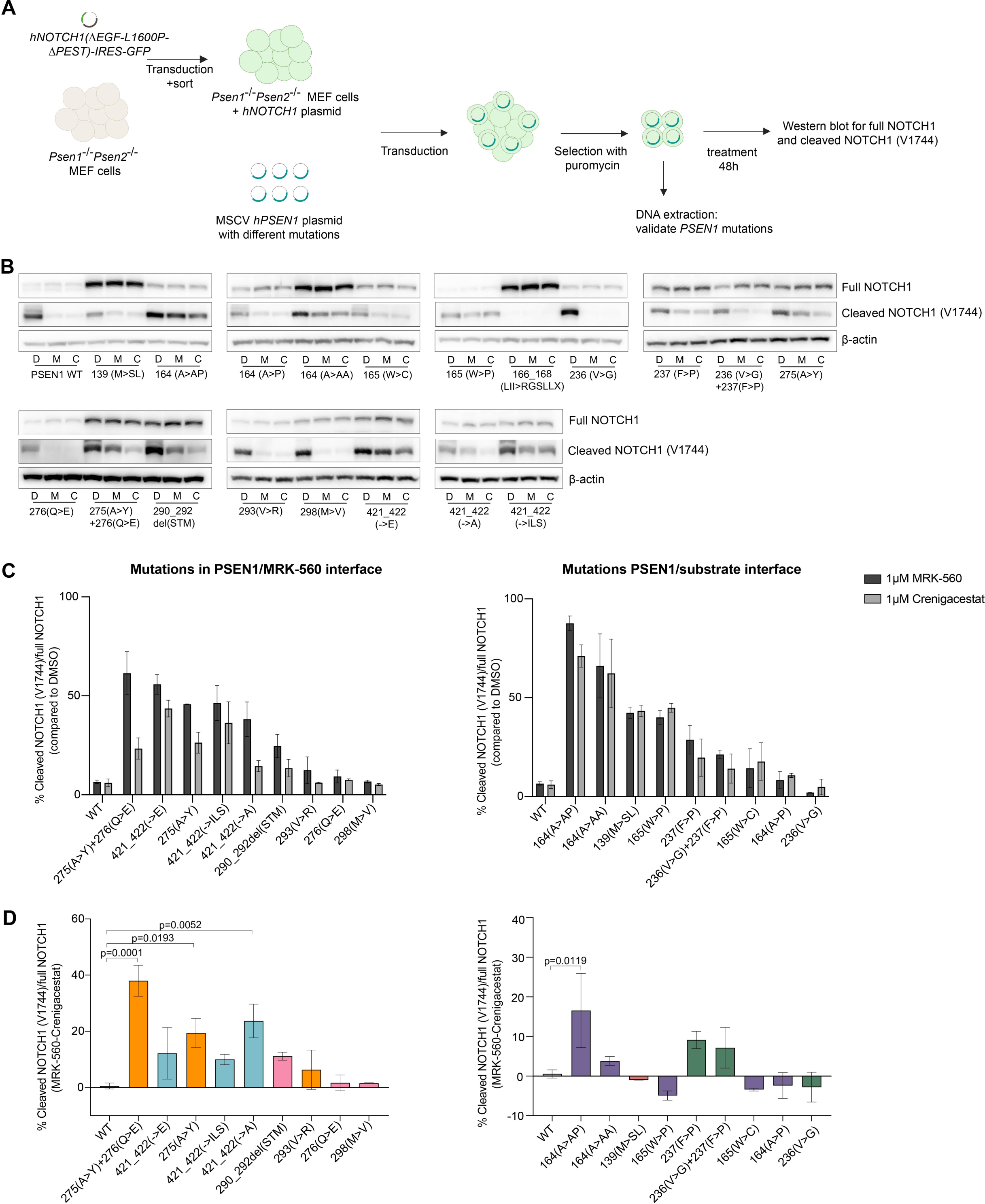
Validation of individual mutations in mouse embryonic fibroblasts identifies the most important amino acid alterations responsible for MRK-560 resistance. **(A)** Experimental set-up of screening of human PSEN1 mutants in *Psen1*^-/-^ *Psen2*^-/-^ mouse embryonic fibroblasts (MEF). **(B)** The MEFs expressing different PSEN1 mutants were treated with DMSO (D), 1 μM MRK-560 (M) or 1 μM crenigacestat (C) for 48 hours and we performed western blotting to detect full NOTCH1 and cleaved NOTCH1(V1744). (**C**) The graphs show the percentage of cleaved NOTCH1 (V1744) normalized to full NOTCH1 for each PSEN1 mutant. High percentage of the ratio cleaved NOTCH1/full NOTCH1 corresponds to high resistance towards the inhibitor. **(D)** The graphs show the difference in percentage of cleaved NOTCH1/full NOTCH1 ratio upon treatment with MRK-560 versus crenigacestat. A high difference indicates more resistance to MRK-560 compared to crenigacestat. Panel C and D show mean and standard deviation of 2 replicates. Statistical differences were obtained with ANOVA Dunnett’s multiple comparison test.

Next, we treated each MEF cell line for 48 hours with DMSO, MRK-560 or a broad-spectrum GSI and determined full and cleaved NOTCH1 (Val1744) (**Fig. 2B-D**). High levels of cleaved NOTCH1 compared to full NOTCH1 after treatment with GSI was indicative of resistance. In cells with wild type PSEN1, NOTCH1 cleavage was effectively inhibited by MRK-560. In contrast, we observed that for the majority of PSEN1 mutants NOTCH1 was still cleaved in the presence MRK-560 treatment, except for mutations 293(V>R), 276(Q>E), 298(M>V), 164(A>P) and 236(V>G) (**Fig. 2C**). The most resistant mutations with high levels of cleaved NOTCH1/full NOTCH1 upon MRK-560 treatment were 275(A>Y) + 276(Q>E), 421_422(->E), 275(A>Y), 421_422(->ILS), 421_422(->A), 164(A>AP), 164(A>AA), 139(M>SL) and 165(W>P). Furthermore, we calculated the differential effect of the PSEN1-selective GSI (MRK-560) and the broad-spectrum GSI (crenigacestat). We noticed that almost all mutations localized in the PSEN1/MRK-560 interface had higher levels of cleaved NOTCH1/full NOTCH1 when treated with MRK-560, compared to crenigacestat (**Fig. 2D**). This also corresponds to our findings from our CRISPR-Cas9 T-ALL screening in **Fig. 1D-E** where cells transduced with sgRNA pools against exon 8 and 12 were most resistant to MRK-560 compared to pan-GSI.

### Alterations in amino acids 275, 276, 421 and 422 of PSEN1 are responsible for selective resistance to MRK-560 in T-ALL cell lines

To validate the resistance mutations obtained from the screen, we used CRISPR genome editing to generate various mutations in the genome of 3 T-ALL cell lines. We designed sgRNAs and homology directed repair (HDR) oligos for generating the following resistant mutations: 275(A>Y), 275(A>Y)+276(Q>E) and 421_422(->ILS) (**Table 1**). We introduced these mutations in DND-41, HPB-ALL and RPMI-8402, three T-ALL cell lines with a mutation in the NOTCH1 receptor and known to be dependent on NOTCH1 signaling for survival. The cell lines already expressed Cas9 and following electroporation with sgRNA and template, we treated the cells with MRK-560 for 3 weeks to select for resistant cells (**Fig. 3A, Fig. EV3A**). Subsequent Sanger sequencing confirmed the presence of the mutations in the majority of the cells (**Fig. EV3B**).

**Figure 3.**
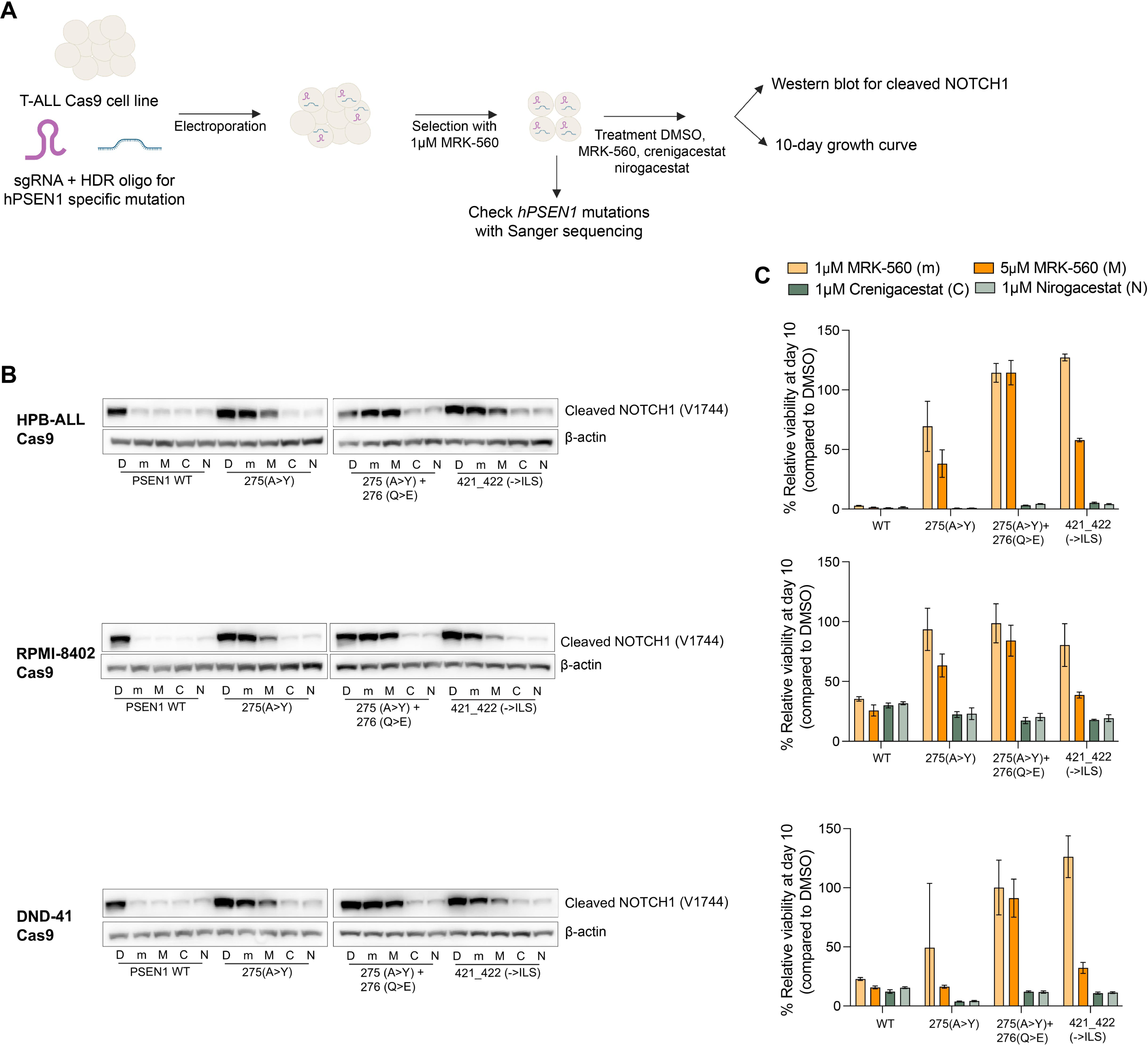
Alterations in amino acids 275, 276, 421 and 422 of PSEN1 results in selective resistance to MRK-560 in T-ALL cell lines. **(A)** Experimental set-up to generate specific *PSEN1* resistance mutations in the genome of DND-41, HPB-ALL, RPMI-8402 cell lines using CRISPR genome editing. The mutations were confirmed with Sanger sequencing and cells were treated with DMSO (D), 1 μM MRK-560 (m), 5 μM MRK-560 (M), 1 μM crenigacestat (C) or 1 μM nirogacestat (N) for 48 hours to detect NOTCH1 cleavage levels (panel B) or for 10 days to evaluate the growth (panel C). **(B)** Protein expression levels of cleaved NOTCH1(V1744) in T-ALL Cas9 cell lines with a specific mutation. **(C)** Relative viability percentages (normalized to DMSO) at day 10 of long-term treatment experiments with each drug condition in the different cell lines. The percentages in panel C show mean and standard deviation of 3 replicates.

These cells were treated with DMSO, MRK-560, crenigacestat and nirogacestat (**Fig. 3B-C**) to assess MRK-560 resistance and the difference in resistance levels between MRK-560 and the pan-GSIs. All three mutations resulted in sustained NOTCH1 cleavage upon treatment with MRK-560 compared to pan-GSIs (**Fig. 3B**). This sustained cleavage of NOTCH1 was also associated with proliferation and survival of the cells during treatment of 10 days (**Fig. 3C, Fig. EV4**). Both experiments validated that mutations at amino acids 275, 276, 421 and 422 conferred resistance to MRK-560 and not to broad-spectrum GSIs.

### Identification of two types of resistance mutations located at the enzyme-drug interface or enzyme-substrate interface

To gain insight in how these mutations confer resistance to MRK-560, we analyzed the secondary and 3-D structures of human PSEN1 together with the position of the resistance mutations obtained in **Fig. 1** (**Fig. 4**). By analyzing the secondary structure, we localized the position of the resistance mutations in transmembrane domain (TMD) 2, 3, 5 and 8 (mutations around amino acids 139, 164, 165, 236, 237, 421, 422). Other mutations found at amino acids 275, 276, 290, 293 and 298 are situated in the cytosolic loop of PSEN1 (**Fig. 4A**). Recently, high resolution cryo-EM structures showed how MRK-560 binds to PSEN1 in the vicinity of the active site (**Fig. 4B**, left panel).^28^ Specifically, the inhibitor binds within the TMD 6 (amino acids 249-272), the intracellular TMD 6a (266-276) and following loop-2 (277-286), the β-sheet 2 that precedes TMD 7 (380-382) and the cytosolic sides of TMD 8 and TMD 9 in PSEN1 (**Fig. 4B**, top panel). MRK-560 is stabilized by hydrogen bonding interactions with the catalytic Asp385 (TMD 7), and the backbones of Leu432 (before the Pro433-Ala433-Leu434 or PAL motif between TMD 8 and 9) and Leu282 (loop-2 between TMD 6a and TMD 7) (shown in pink in **Fig. 4B**), as well as by extensive van der Waals contacts with residues in the binding pocket (**Table 3**).^28^

**Figure 4.**
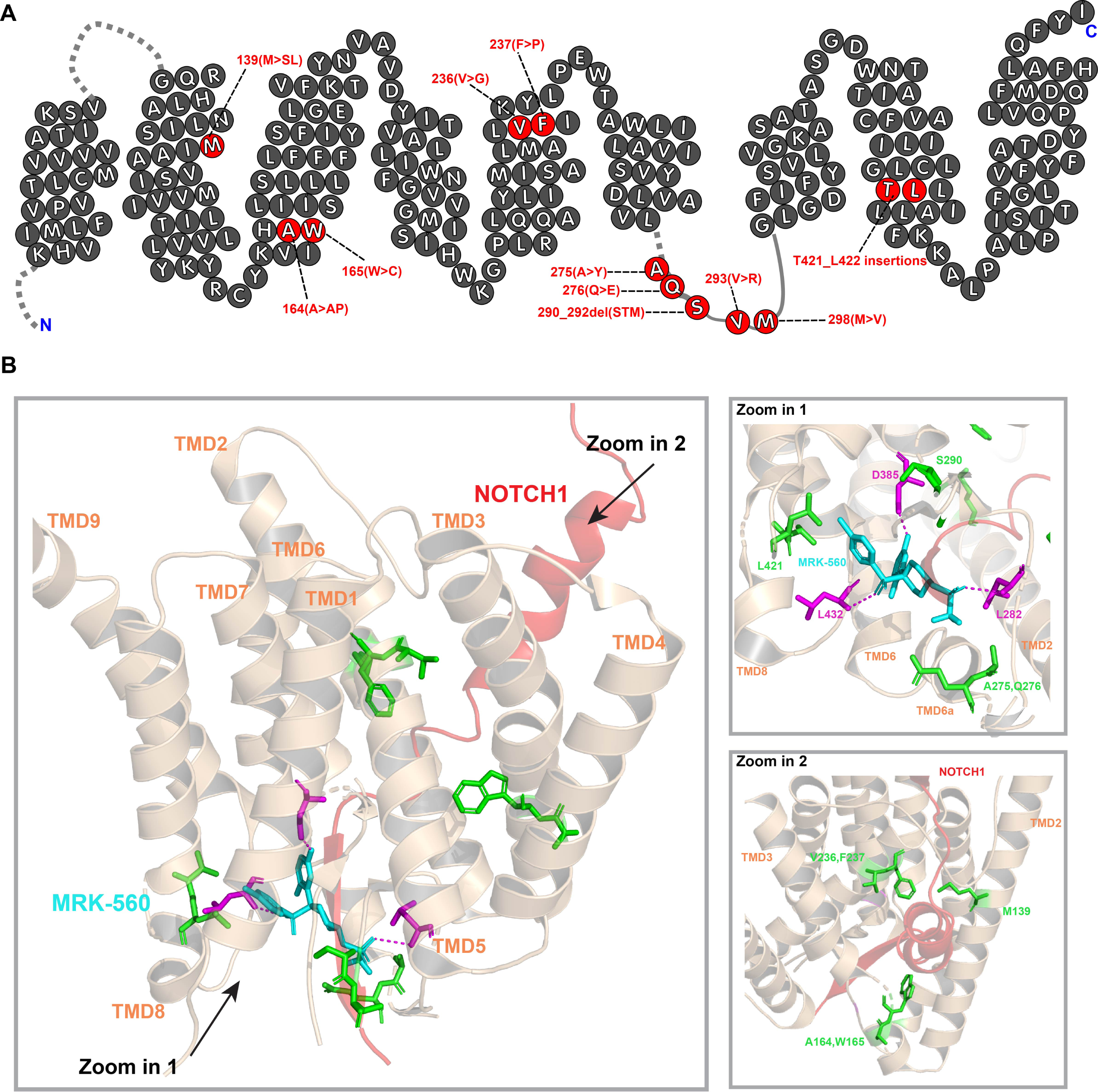
The localization of the resistance mutations on secondary and 3-D PSEN1 protein structures provides insight in potential resistance mechanisms. (**A**) Secondary protein structure of human PSEN1 with the potential resistance mutations highlighted in red. The structure was adapted from Szaruga et al. (2017).^30^ **(B**) 3-D structure of human PSEN1, together with human NOTCH1 (red) and MRK-560 (blue) (Protein Data Bank:7Y5T and 6IDF). Known interaction partners of MRK-560 with PSEN1 are highlighted in pink (hydrogen bounds) and newly identified resistance positions obtained in this study in green. Figures are made with PyMol.

**Table 3.** Overview of altered amino acids of PSEN1 that confer resistance to MRK-560^28^.

| Position | Location | PSEN1/PSEN2 conserved? | Type of interaction |
| --- | --- | --- | --- |
| ***Known direct interaction partners of MRK-560*** |  |  |  |
| Val272 | TMD 6a | Yes | Van der Waals |
| Thr281 | Loop-2 (after TMD 6a) | No | Van der Waals |
| Leu282 | Loop-2 (after TMD 6a) | No | Hydrogen bound |
| Val379 | $\beta$ sheet 2 (Preceding TMD 7) | Yes | Van der Waals |
| Lys380 | $\beta$ sheet 2 (Preceding TMD 7) | Yes | Van der Waals |
| Gly382 | $\beta$ sheet 2 (Preceding TMD 7) | Yes | Van der Waals |
| Asp385 | TMD 7 | Yes | Hydrogen bound |
| Leu418 | TMD 8 | Yes | Van der Waals |
| Thr421 | TMD 8 | Yes | Van der Waals |
| Leu422 | TMD 8 | Yes | Van der Waals |
| Ala431 | PAL motif (Between TMD 8 and 9) | Yes | Van der Waals |
| Leu432 | PAL motif (Between TMD 8 and 9) | Yes | Hydrogen bound |
| Pro433 | PAL motif (Between TMD 8 and 9) | Yes | Van der Waals |
| Ala434 | PAL motif (Between TMD 8 and 9) | Yes | Van der Waals |
| ***Newly identified (in)direct interaction partners of MRK-560*** |  |  |  |
| Ala275 | TMD 6a | Yes | Close to Val272 |
| Tyr276 | TMD 6a | Yes | Close to Val272 |
| Ser290 | Loop-3 (Between TMD 6a and 7) | Yes | Stabilizing β1-β2 sheet |
| Thr291 | Loop-3 (Between TMD 6a and 7) | No | Stabilizing β1-β2 sheet |
| Met292 | Loop-3 (Between TMD 6a and 7) | Yes | Stabilizing β1-β2 sheet |
| Val293 | Loop-3 (Between TMD 6a and 7) | Yes | Stabilizing β1-β2 sheet |
| ***Amino acids important for recognition and stabilization of NOTCH1/APP substrates*** |  |  |  |
| Met139 | TMD 2 | Yes | Unknown interaction |
| Ala164 | TMD 3 | No | Unknown interaction |
| Trp165 | TMD 3 | Yes | Hydrogen bound with Ser169, which forms hydrogen bound with APP/NOTCH1 |
| Phe237 | TMD 5 | Yes | Van der Waals with APP/NOTCH1 |

We asked whether the identified mutations (shown in green in **Fig. 4B**) alter the binding of MRK-560 to PSEN1 by disrupting its binding pocket. Based on the distance of the mutation to MRK-560 in the 3-D structure, we identified mutations situated either directly at the interface between MRK-560 and PSEN1 (**Fig. 4B zoom in 1**), or localized in the PSEN1-NOTCH1 interface (**Fig. 4B zoom in 2**). In the first group, the identified resistance mutations were found at the intracellular TMD 6a (Ala275, Gln276), the cytosolic loop-3 (Ser290-Val293) and in the cytosolic end of TMD 8 (Thr421, Leu422). In the second group, the resistance mutations mapped to TMD 2 (Met139), TMD 3 (Ala164, Trp165) and TMD5 (Val236, Phe237) (**Table 3)**.

### Mutations located at the enzyme-substrate interface change affinities of PSEN1 for substrate and/or MRK-560, leading to GSI resistance

To further unravel the resistance mechanisms of mutations located at the enzyme-substrate interface, we used the MEF cell lines to investigate the cleavage of NOTCH1 by wild type/mutant PSEN1s in the presence of increasing concentrations of MRK-560. Initially, we assessed the baseline levels of cleaved NOTCH1/full NOTCH1 for wild type PSEN1 and mutant PSEN1 cell lines (**Fig. 5A, Fig. EV5A**). Notably, most mutations in the enzyme-drug interface showed significantly higher levels of cleaved NOTCH1/full NOTCH1 compared to wild type PSEN1, suggesting that these *PSEN1* mutations increased the affinity and/or turnover of NOTCH1. In contrast, mutations in the enzyme-substrate interface have similar or lower levels of NOTCH1 cleavage, indicating no changes or reduced affinity/turnover for NOTCH1. Subsequently, we treated selected MEF cell lines with DMSO or increasing doses of MRK-560 for 24 hours to generate dose response curves for the levels of cleaved NOTCH1 (**Fig. 5B-C, Fig. EV5B**). These curves provide information on the relative affinities of PSEN1 for MRK-560, quantified in IC50 values. We observed higher IC50 values (> 200 fold compared to wild type) for mutations 164(A>AP) and 275(A>Y), which were associated with lower affinities for MRK-560.

**Figure 5.**
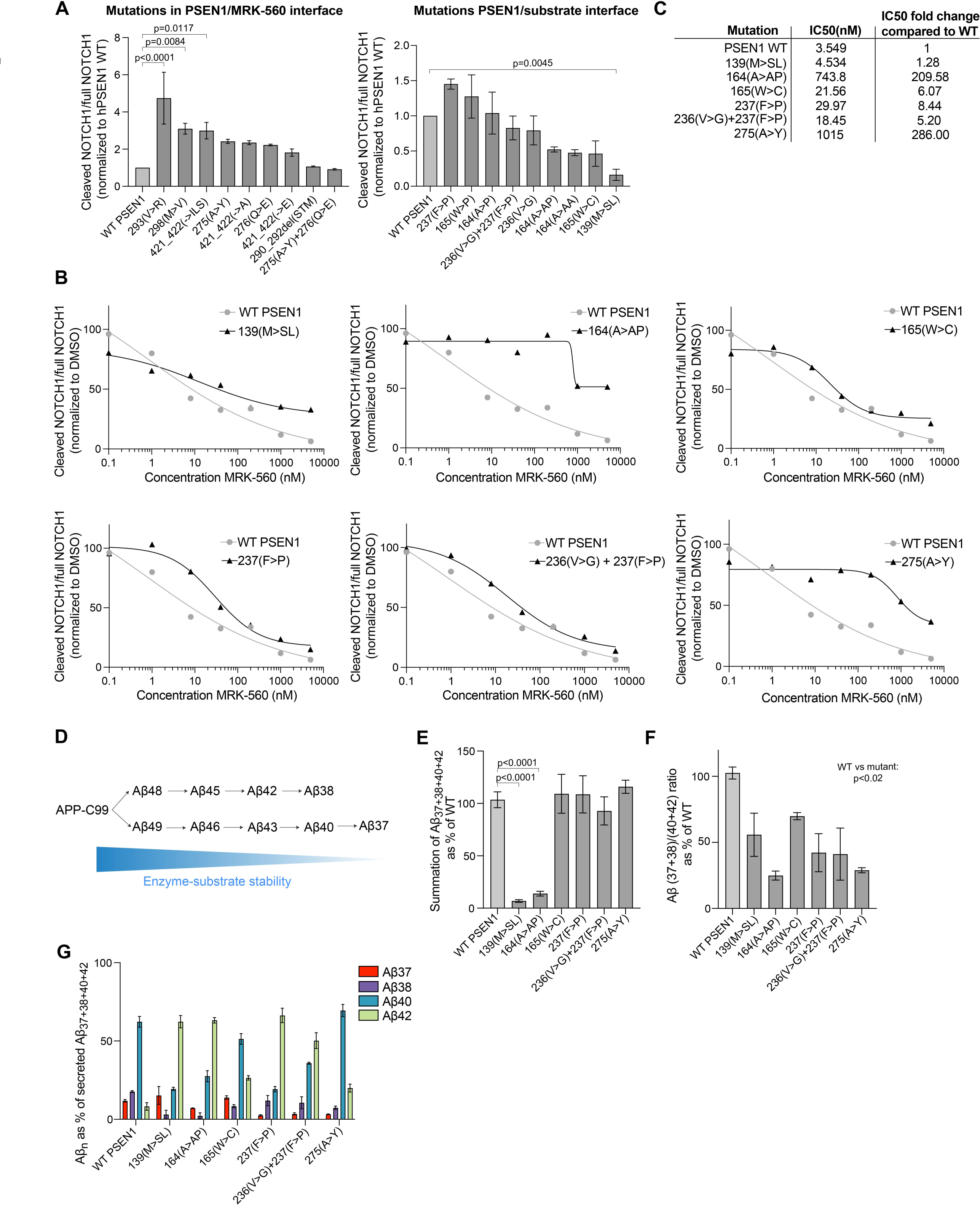
*PSEN1* mutations located at the enzyme-substrate interface lower substrate kinetics. **(A)** Protein levels of cleaved NOTCH1(Val1744)/full NOTCH1 in MEF cell lines expressing human NOTCH1(ΔEGF-L1600P-ΔPEST) with wild type (WT) or mutated human PSEN1. The cleaved/full NOTCH1 ratio of each mutant was normalized to the ratio of PSEN1 WT. **(B)** Protein levels of cleaved NOTCH1/full NOTCH1 of MEF cell lines + NOTCH1 (ΔEGF-L1600P-ΔPEST) with WT or mutated PSEN1 after treatment for 24 hours with DMSO or increasing concentrations of MRK-560. These ratios were normalized to the ratio after DMSO treatment. **(C)** IC50 values derived from dose response curves obtained in panel B. **(D)** Sequential cleavages of the amyloid precursor protein (APP)-C99 by the γ-secretase complex. Stability of the enzyme-substrate decreases with each cleavage event.^29^ **(E)** Summation of Aβ37+Aβ38+Aβ40+Aβ42 peptide levels which are normalized to WT PSEN1. **(F)** Aβ(37+38)/(40+42) ratios normalized to PSEN1 WT. **(G)** Quantification of Aβ peptides (37,38,40,42), which were secreted by MEF cells in supernatant after transduction with APP-C99 virus by multiplex MSD ELISA. We showed the proportion of each Aβ peptide. Panel A shows mean and standard deviation of 2 replicates, while panels F,G and H include 3 replicates. Statistical differences were obtained with ANOVA Dunnett’s multiple comparison test.

We then investigated the effects of *PSEN1* mutations, located at the enzyme-substrate interface, on the processing of amyloid precursor protein (APP) by quantifying the levels of Aβ37, Aβ38, Aβ40, Aβ42 peptides. APP is another substrate of the γ-secretase complex that undergoes sequential cleavages by PSEN1 (**Fig. 5D**).^29^ Interestingly, pathogenic mutations in *PSEN1* that cause early-onset Alzheimer’s disease reduce the stability of PSEN1-APP/Aβ interactions during the sequential proteolysis, thereby leading to the premature release of longer Aβ fragments (Aβ42 and Aβ43).^30^ Since the structures of APP and NOTCH1 bound to PSEN1/γ-secretase show a similar overall enzyme-substrate conformation, we reasoned that alterations in the length of Aβ fragments generated from APP by resistance mutations in *PSEN1*, would inform on possible (de)stabilization effects of enzyme/substrate (APP/NOTCH1) interactions.^31,32^

We created new MEF cell lines with PSEN1 wild type or mutant forms, which all rescued the activity of the γ-secretase complex (**Fig. EV5C**). These cell lines were transduced with APP-C99 adenovirus and secreted Aβ peptides were measured in the conditioned media with multiplex MSD ELISA. First, we estimated the global endopeptidase activity of the γ-secretase complex for APP using as a proxy the total Aβ37 + Aβ38 + Aβ40 + Aβ42 levels. We observed lower global activity for mutations 139(M>SL) and 164(A>AP), indicating a lower affinity and/or turnover for APP (**Fig. 5E**). The same *PSEN1* mutations also decreased NOTCH1 processing (**Fig. 5A**). The 165(W>C), 237(F>P), 236(V>G)+237(F>P) and 275(A>Y) mutations did not result in a significant change in total Aβ production. Similarly, these mutations caused minor changes in cleaved NOTCH1 levels. In order to assess the γ-secretase processivity, we delved deeper into the proportion of each Aβ peptide for the different cell lines. The Aβ profiles indicated a shift towards the production of longer Aβ fragments (Aβ40 and Aβ42) compared to wild type PSEN1 for all mutants (**Fig. 5G**). To estimate the efficiency of the sequential γ-secretase cleavage, we calculated the ratio of Aβ (37+38)/(40+42). We found a significant decrease in γ-secretase processivity for all the mutants compared to wild type PSEN1 (**Fig. 5F**).

## Discussion

PSEN1-selective GSIs are a potential new treatment option for NOTCH1 mutant T-ALL cases. These selective GSIs have shown efficacy in T-ALL cell lines in vitro as well as in PDX models in vivo and due to the selectivity treatment-related toxicities are limited. Despite these promising pre-clinical findings, it remains unknown whether mutations in *PSEN1* could lead to resistance against PSEN1-selective GSIs in T-ALL. Therefore, we conducted a CRISPR screen in a T-ALL cell line to introduce random mutations in *PSEN1* and investigate their potential to induce resistance to MRK-560 treatment.

In our initial CRISPR-Cas9 screening in the T-ALL RPMI-8402 Cas9 cell line, we found that mutations in exon 6 of *hPSEN1* (164(A>AP), 165(W>C)) resulted in high resistance to both the PSEN1-selective inhibitor MRK-560 and other pan-GSIs. Conversely, mutations in exon 8 (275(A>Y),276(Q>E)) and exon 12 (421_422 insertions) conferred pronounced resistance specifically towards MRK-560. Moreover, we validated that mutations in amino acids 275, 276, 421 and 422 were indeed responsible for selective resistance to MRK-560 in mouse embryonic fibroblasts and T-ALL cell lines.

In order to gain deeper insights into the resistance mechanisms and the PSEN1-selectivity of all *PSEN1* mutations we discovered, we used the cryo-EM structures of the MRK-560 binding pocket to explain the observed resistance phenotypes (**Table 3**).^28^ Firstly, mutations located at the enzyme-drug interface, resulting in selective resistance to the PSEN1-selective GSI, likely interfere directly with the recognition and/or binding of MRK-560. Insertions around amino acids Thr421 and Leu422 disrupted the cytosolic end of TMD 8. This region (421-422) is conserved in PSEN1 and PSEN2 and is involved in MRK-560 recognition via van der Waals interactions with the inhibitor’s chlorophenyl group.^28,33^ Therefore, insertions between amino acids 421 and 422 likely disturb key interactions between Thr421-Leu422 and MRK-560. We also hypothesize that insertions between 421 and 422 might shift the subsequent residues, including Leu432 and the critical PAL-motif. Leu432 is known to form a hydrogen bound with MRK-560 and the mutation-driven shift can also disrupt this key interaction. Notably, Leu432 is located further away from other broad-spectrum GSIs, such as avagacestat, which could explain the selective resistance towards MRK-560 in this type of mutations.

Substitutions affecting the intracellular TMD 6a (275(A>Y) + 276(Q>E), 275(A>Y)) also conferred a high degree of resistance to MRK-560. In the intracellular TMD 6a helix, Val272 interacts with the trifluoromethanesulfonamide (TFMS) moiety of MRK-560.^28^ The resistance conferred by the mutations indicates that introduction of a larger (A > Y) and/or a negatively charged (Q > E) residue in the vicinity (positions 275-276, respectively) disrupts the interactions with the inhibitor’s TFMS moiety. These mutations alter recognition, but not selectivity, as this region is conserved between PSEN1 and PSEN2. Intriguingly, the larger substitution at position 275 (A>Y) displayed differential effects on the response to MRK-560 and crenigacestat/nirogacestat, implying a differential involvement of the intracellular TMD 6a helix in the binding and recognition of these inhibitors. Structural data for other broad-spectrum GSIs avagacestat and semagacestat indeed shows a larger distance between PSEN1 Ala275/Gln276 and the drug, compared to MRK-560.^28,34^ In addition, Guo *et al.* confirmed that loop-2 (located directly downstream of TMD 6a) is closer located to MRK-560, further highlighting the importance of both TMD 6a and loop-2 in the subtype specific character of MRK-560.^28^

In addition to a direct interference with MRK-560 recognition, mutations at amino acids 275, 421 and 422 may also have an indirect effect. The MRK-560 binding site within PSEN1 is common to several GSI inhibitors and partially overlaps with the binding site of the substrate.^34^ Therefore, we investigated if this type of mutations could also affect substrate processing. Our experiments showed that mutations of amino acids 275, 421 and 422 resulted in slightly higher NOTCH1 cleavage compared to wild type PSEN1 (**Fig. 5A)**. This implies an increased affinity and/or turnover of NOTCH1, which could hamper the binding of MRK-560 due to competition, given that both inhibitor and substrate bind to an overlapping site in PSEN1. Indeed, mutation 275(A>Y) in *PSEN1* resulted in a higher IC50 value for MRK-560, when assessing cleaved NOTCH1 (**Fig. 5C**). This demonstrates a lower drug affinity. A similar inhibitor-substrate effect has been shown for the γ-secretase inhibitors DAPT and semagacestat APP by Koch *et al.*^29^ DAPT and semagacestat, which bind to the common GSI site, act as high-affinity competitors to the substrate and mutations that stabilize enzyme-substrate interactions (thus favoring the interaction with the substrate) shifted the IC50 values for DAPT and semagacestat towards higher concentrations.^29^

Moreover, we identified deletion of Ser290-Thr291-Met292 in hPSEN1, located in the proximity of the PSEN1/MRK-560 interface, that also resulted in resistance. This 290_292del(STM) mutation affects the region immediately following the intracellular β1 structure, which is part of a large intracellular loop-3. It is known that the common GSI binding pocket, including the MRK-560 binding pocket, is stabilized by the β1-β2 sheet structure.^28,31^ The Ser290/Thr291 amino acids are involved in hydrogen bonding with residues preceding the β2 structure and are likely to contribute to the stability of the β1-β2 structure. Deletion of the STM in PSEN1 (SAM in PSEN2) fuses the C-terminus of the β-sheet 1 with relatively larger and hydrophobic residues (VWL) which cannot engage in H-bonding interactions stabilizing the β1-β2 structure. Swapping loop-3 between PSEN1 and PSEN2 had no detectable effect on the sensitivity to MRK-560, suggesting that the resistance conferred by the 290_292del(STM) mutation may be due to the loss of two H-bonds that contribute to the stability of the β1-β2 structure and thus to general GSI binding.^28^ In addition, a steric effect of the larger side chains of the W residue, occupying the position of the ST in the mutant 290_292del(STM), may contribute to the resistance. In our experiments, we did not find a large difference in resistance to MRK-560 or crenigacestat/nirogacestat when this mutation was present, further supporting that the β1-β2 structure contributes to the general GSI binding pocket.

The second group of resistance mutations map to TMD 2, 3 and 5. These TMDs are known to contribute to the recognition and binding of the substrates.^31,35^ We hypothesize that 139(M>SL), 164(A>AP), 165 (W<C) and 237 (F>P) mutations probably affect the overall folding of PSEN1 and perturb the interface of PSEN1 with the substrate and/or drug (**Fig 4B, bottom right panel**). To challenge this hypothesis, we investigated the substrate and drug affinities in the presence of these *PSEN1* mutations (**Fig. 5)**. Mutations 139(M>SL), 164(A>AP) and 165(W>C) resulted in a reduced level of NOTCH1 cleavage compared to wild type PSEN1 in the absence of GSI, indicating a decrease in affinity and/or turnover of NOTCH1. Given the high similarity in the binding conformation between NOTCH1 and APP to PSEN1, we also evaluated the processing of APP. Specifically, we assessed potential mutation-driven effects on the destabilization of γ-secretase and APP/Aβ interactions, which is reflected in a relative increased production of longer Aβ fragments. ELISA-based analysis (**Fig. 5D-G**) showed that all resistance mutations increased the production of longer Aβ fragments, arguing in favour of mutation-induced destabilization of enzyme-substrate interactions. Data showing that mutations do not lower total Aβ peptide levels indicate that the destabilization is relatively mild and mainly affect the interaction with Aβ peptides (shown to be less stable than the one with the initial APP substrate).^30^ In addition, the MRK-560 dose-response curves and derived IC50 values (**Fig. 5B-C**) showed that the evaluated mutations reduce the affinity to MRK-560, compared to wild type PSEN1.

Therefore, we hypothesized that the 139(M>SL), 164(A>AP), 165(W>C) and 237(F>P) mutations lower the stability of the interactions between PSEN1 and the substrate (NOTCH1/APP). However, a greater detrimental impact on the affinity to MRK-560, demonstrated by the IC50 values, probably leads to resistance to GSIs. Interestingly, we observed a pronounced effect on drug binding for the mutation 164(A>AP), suggesting an additional effect. Recently, Serneels *et al.* found that residue Leu172 plays a crucial role in facilitating the entry of PSEN1-selective inhibitors in the PSEN1 pocket.^33^ Their findings revealed that substitution of leucine with alanine, resulting in a wider entrance channel, enhanced the entry of MRK-560 to a greater extent compared to pan-GSIs. Given that our mutation 164(A>AP) is proximal of Leu172, we can hypothesize that introducing a bulkier pyrrolidine moiety will have higher impact on MRK-560 entry compared to pan-GSIs, thus leading to higher MRK-560 resistance levels.

In conclusion, our findings revealed that treatment with MRK-560 in T-ALL cell lines can induce drug resistance, that is attributed to mutations in *PSEN1*. We identified three potential resistance mechanisms that rescue the NOTCH1 signaling in the presence of GSIs. Firstly, amino acids located in the enzyme-drug interface directly disrupt the binding of GSIs. We showed that amino acids Ala275, Thr421 and Leu422 are important for specific binding of the PSEN1-selective drug, highlighting the importance of these regions for PSEN1 selectivity. Additionally, we also observed *PSEN1* mutations at the enzyme-substrate interface. We hypothesize that these mutations alter the relative binding affinities for drug and substrate, leading to more NOTCH1 cleavage. These mutations gave resistance to both types of γ-secretase inhibitors. Lastly, we observed a strong resistance mutation in Ala164, located in the enzyme-substrate interface, that could potentially hinder the entrance of MRK-560 only, emphasizing its importance for PSEN1 selectivity.

Our data illustrate how T-ALL could become resistant to PSEN1-selective GSI and provides insight in the various resistance mechanisms. These data can help to design new PSEN1-selective GSIs to target NOTCH1 activation and to overcome resistance induced by MRK-560.

## Author contributions

CV: study design, methodology, performing experiments, data analysis, writing original draft; SD: analyzing bulk DNA sequencing data, generation fish plot, writing original draft; KJ, LB, SGF: performing experiments; HS: writing original draft, supervision; JC, LCG: study design, methodology, supervision, writing original draft

## Data sharing statement

The data is available by contacting the corresponding authors.

## Acknowledgments

This project was funded by the Belgian Foundation Against Cancer (2020-100 to JC) and KU Leuven (grant C14/18/104 to JC and HS). Bulk DNA sequencing was performed in collaboration with the VIB nucleomics core. We also want to thank Burcu Özcan for providing general protocols for culturing mouse embryonic fibroblasts.

## Expanded view figure legends

**Figure EV1.**
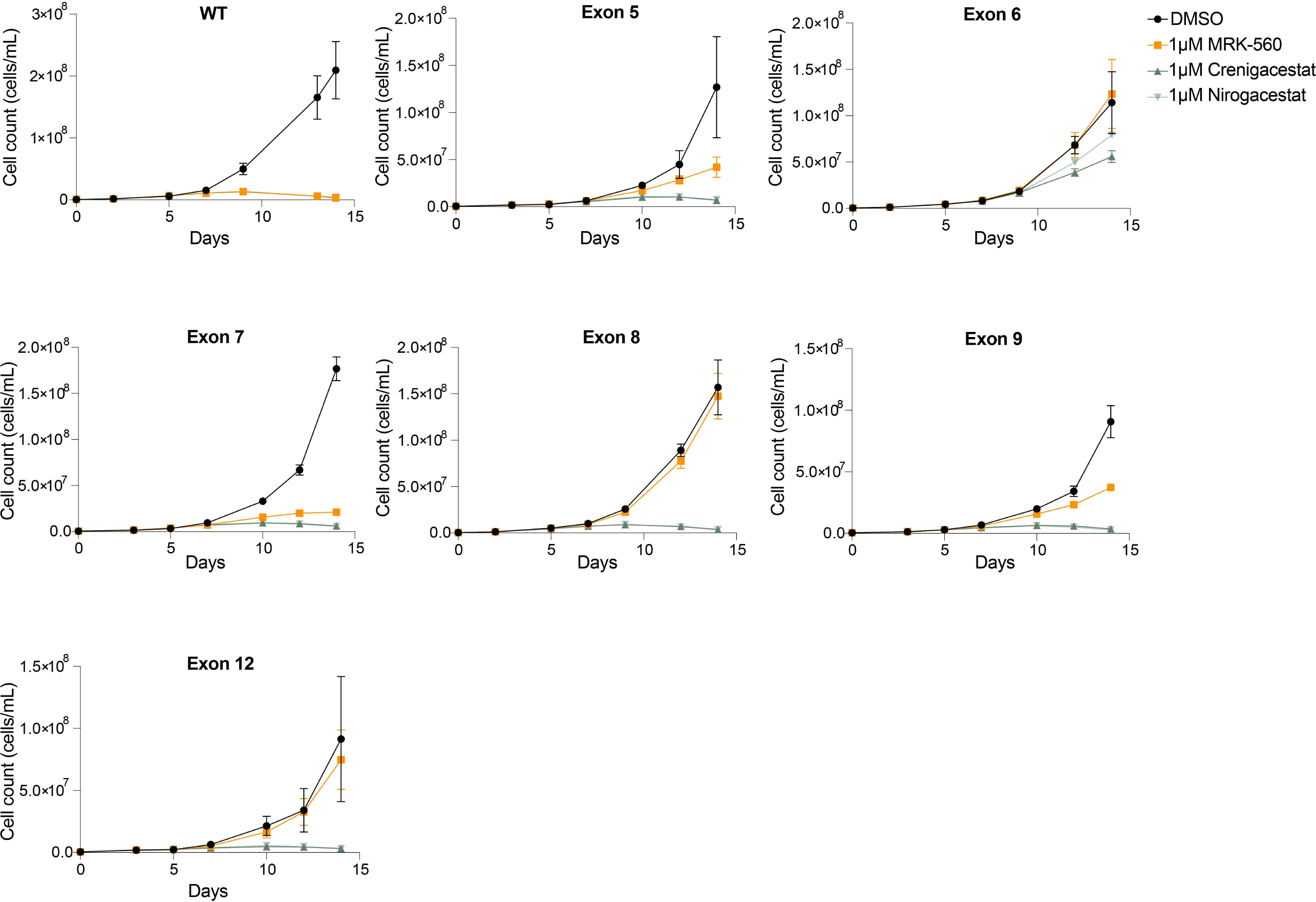
CRISPR-Cas9 mutations in exon 6, 8 and 12 of *hPSEN1* show the highest resistance to MRK-560. Growth curves of resistance cells obtained from RPMI-8402 Cas9 screening from Fig. 1A with DMSO, 1 μM MRK-560, 1 μM crenigacestat and 1 μM nirogacestat for 2 weeks. All growth curves contain mean and standard deviation (error bars) of 3 replicates.

**Figure EV2.**
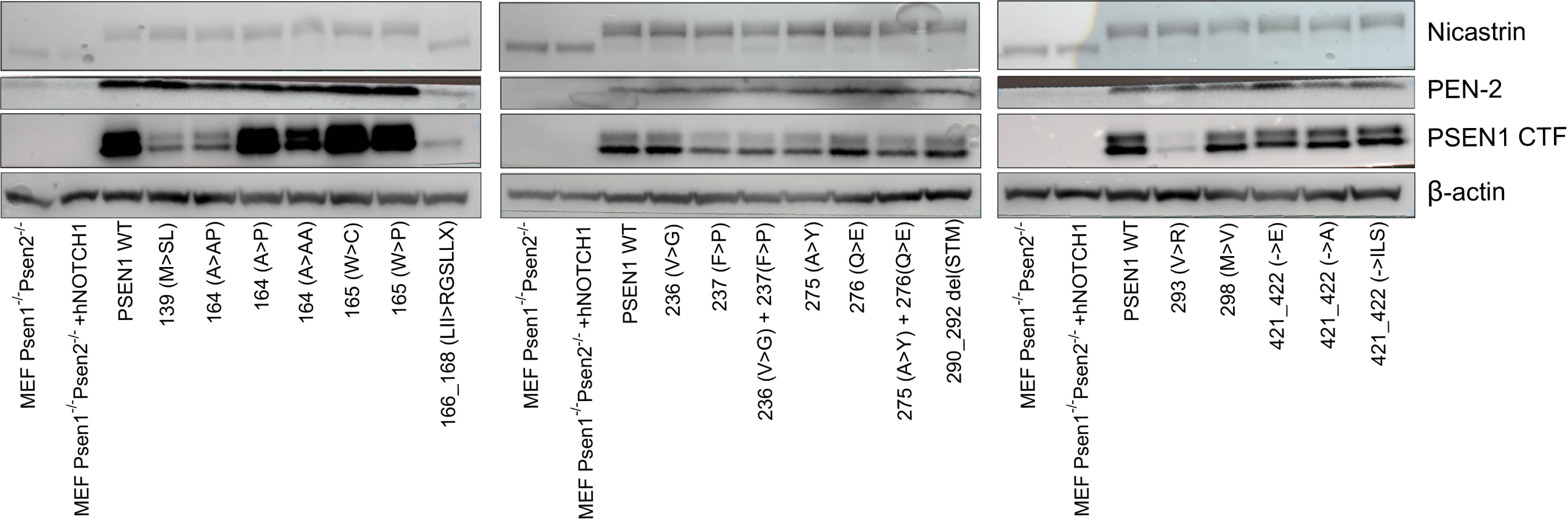
Almost all mutant hPSEN1 are incorporated in the γ-secretase complex of mouse embryonic fibroblasts, rescuing its activity. Western blot for nicastrin (96kDa), PEN-2 (14kDa), PSEN1 CTFs (20-23kDa) and β-actin (43kDa) for Psen1^-/-^ Psen2^-/-^ mouse embryonic fibroblasts (MEF), for Psen1^-/-^ Psen2^-/-^ MEF + hNOTCH1(ΔEGF-L1600P-ΔPEST) and for Psen1^-/-^ Psen2^-/-^ MEF + hNOTCH1(ΔEGF, L1600P, ΔPEST) + hPSEN1 wild type (WT) or hPSEN1 mutant.

**Figure EV3.**
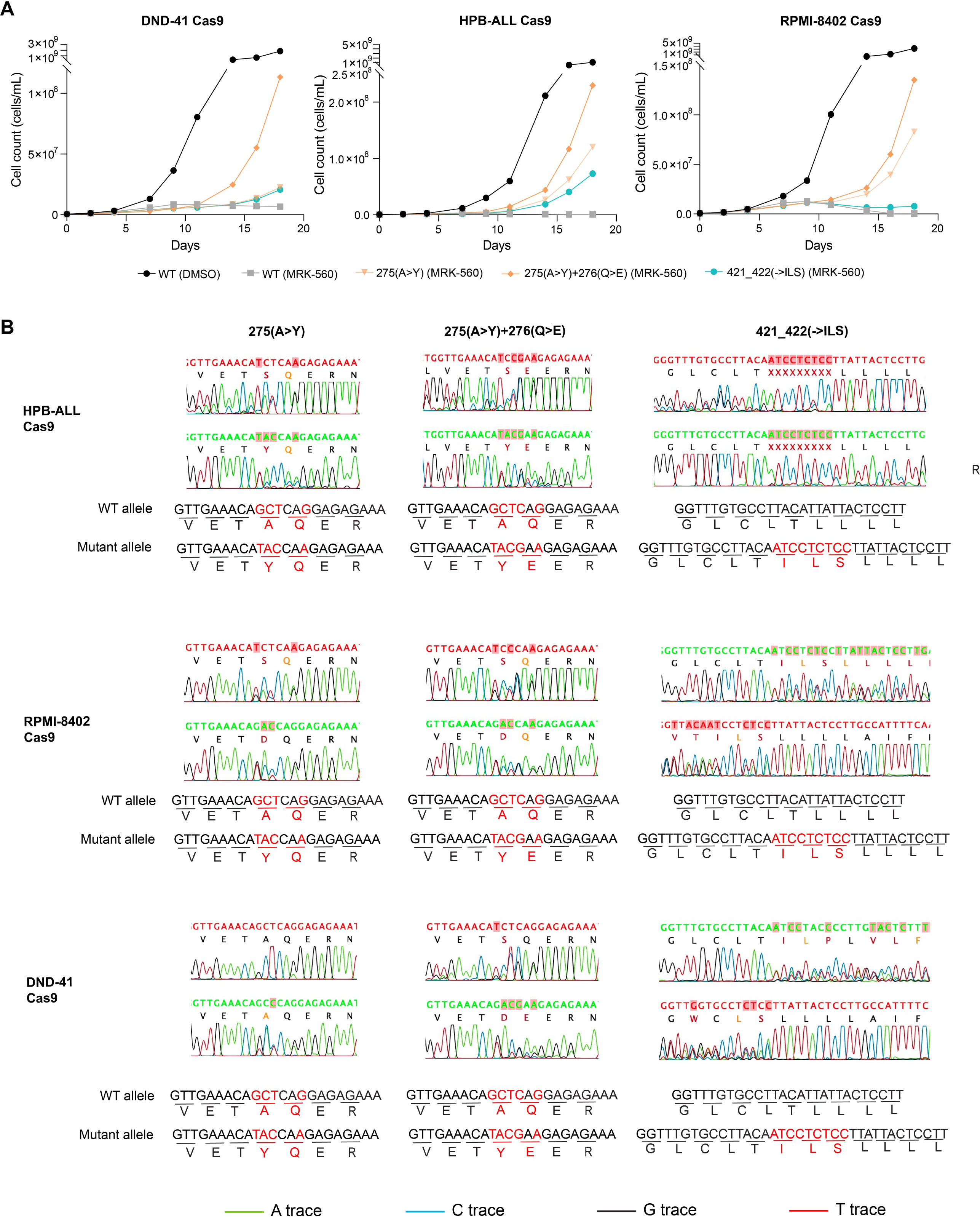
T-ALL Cas9 cell lines obtain resistance to MRK-560 after introduction of a new resistant mutation in their genome with HDR CRISPR. **A)** T-ALL Cas9 cell lines were electroporated with a specific combination of a sgRNA and HDR oligo to introduce a certain mutation in *hPSEN1*. These electroporated cells were subsequently treated with 1 μM MRK-560 for 3 weeks to select for the cells with the desired mutation. **B)** Sanger sequencing data of T-ALL cas9 cell lines after HDR CRISPR and after selection with MRK-560. HDR: Homology Directed Repair.

**Figure EV4.**
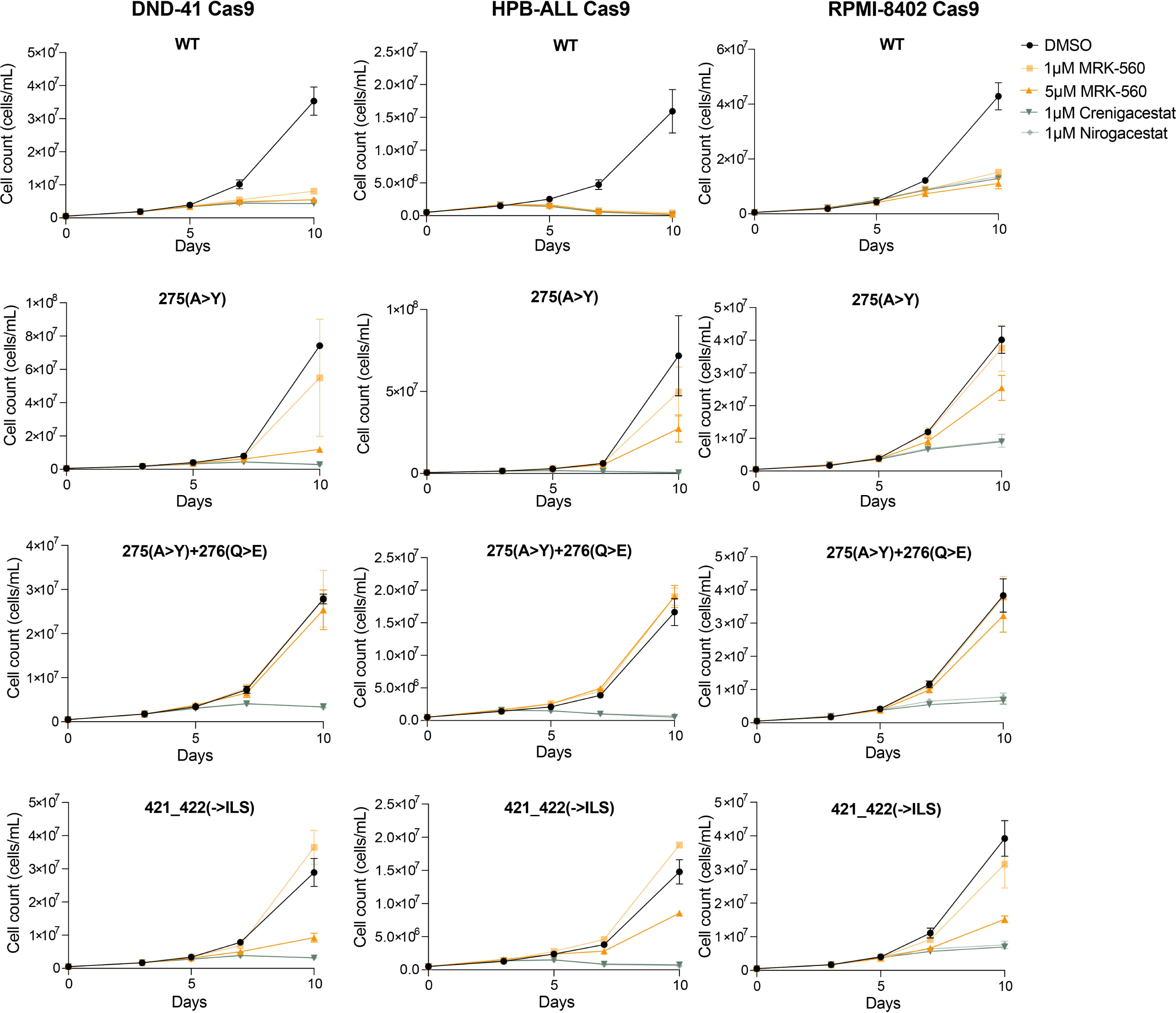
Mutations in amino acids 275, 276, 421 and 422 of hPSEN1 in T-ALL cell lines result in selective resistance to MRK-560. T-ALL Cas9 cell lines with a specific mutation in *hPSEN1* obtained via HDR CRISPR were treated for 10 days with DMSO, 1 μM MRK-560, 5 μM MRK-560, 1 μM crenigacestat or 1 μM nirogacestat. All growth curves contain mean and standard deviation (error bars) of 3 replicates.

**Figure EV5.**
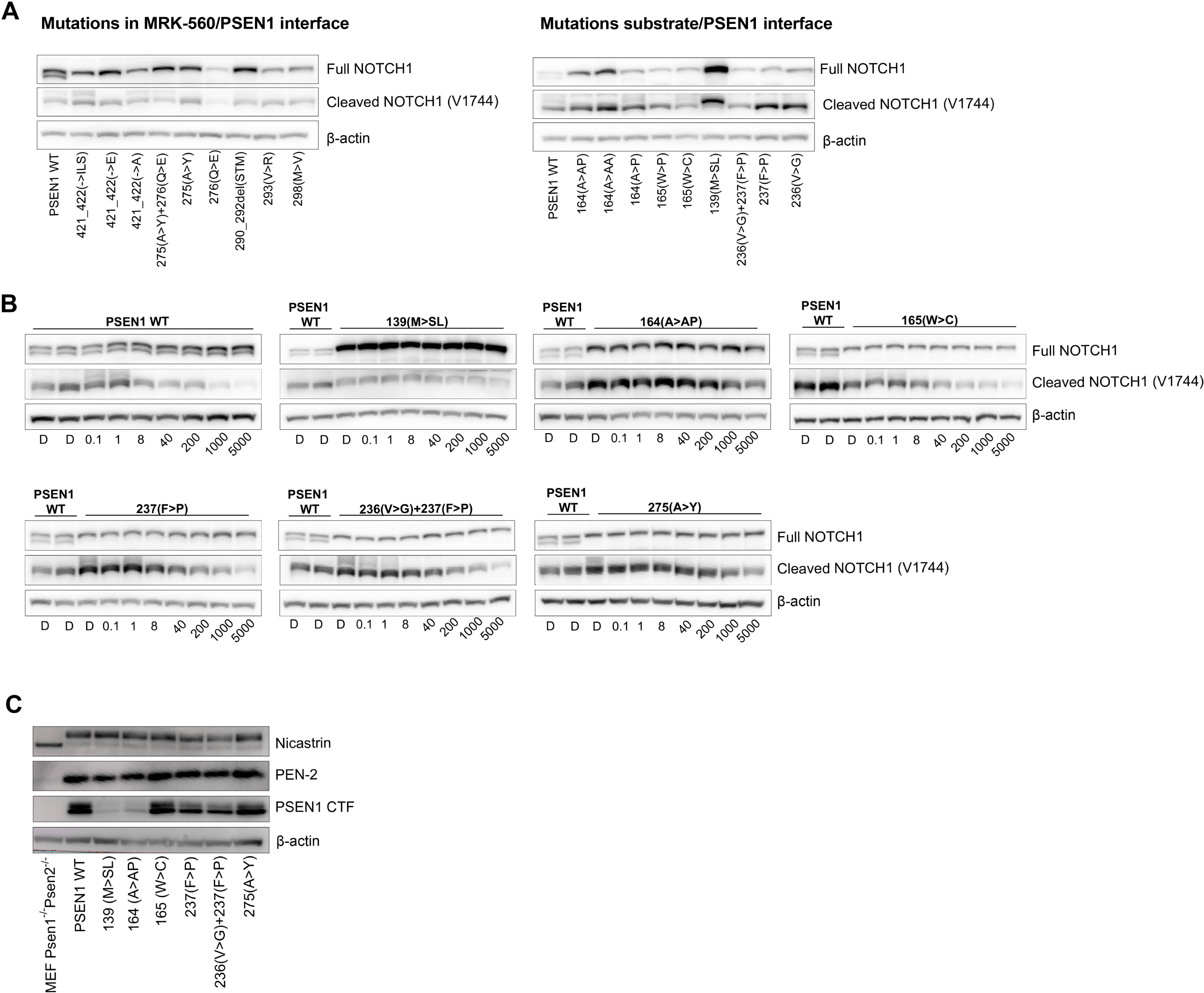
The ratio cleaved NOTCH1/full NOTCH1 determines the substrate affinity and/or turnover. **(A)** Western blot for full NOTCH1, cleaved NOTCH1 (Val1744) and β-actin of Psen1^-/-^ Psen2^-/-^ mouse embryonic fibroblasts (MEF) cells + hNOTCH1 (ΔEGF-L1600P-ΔPEST) transduced with wild type (WT) or mutated *hPSEN1* construct. **(B)** Western blot for full NOTCH1, cleaved NOTCH1 and β-actin of MEF cell lines + hNOTCH1 (ΔEGF-L1600P-ΔPEST) with WT or mutated PSEN1 after treatment for 24h with DMSO (D) or 0.1 nM, 1n M, 8 nM, 40 nM, 200 nM, 1000 nM, 5000 nM MRK-560. **C)** Western blot for nicastrin (96kDa), PEN-2 (14kDa), PSEN1 CTFs (20-23kDa) and β-actin (43kDa) for Psen1^-/-^ Psen2^-/-^ mouse embryonic fibroblasts (MEF) and for Psen1^-/-^ Psen2^-/-^ MEF + hPSEN1 wild type (WT) or hPSEN1 mutant.

